# Real-time, label-free assessment of cell fusion dynamics by high-content imaging

**DOI:** 10.64898/2026.04.08.717136

**Authors:** Sandip Shinde, Anshul Bhide, Pratik Rasal, Deepak Modi

## Abstract

Cell–cell fusion is a fundamental biological process underlying diverse physiological and pathological phenomena, yet its quantitative analysis remains methodologically challenging due to its dynamic, heterogeneous, and multistep nature. Existing approaches to assess fusion largely rely on endpoint assays or manual scoring, limiting temporal resolution, scalability, and reproducibility. Here, we present a label-free, high-content live-cell imaging pipeline for real-time quantification of cell fusion dynamics, developed and validated using trophoblast syncytialization as a model system.

The method integrates automated image acquisition with a reproducible, stepwise analysis workflow combining supervised texture-based segmentation, morphology-based measurements, and intensity-independent texture analysis. We define quantitative metrics, including the ratio of total cluster area to the number of detected clusters and cytoplasmic granularity features, that together discriminate bona fide fusion events from non-fusion–related cellular clustering or proliferation. Using canonical pharmacological inducers and inhibitors of fusion, we demonstrate the specificity and sensitivity of these parameters for detecting fusion-associated remodeling over time.

We further demonstrate the scalability of the pipeline through high-throughput screening of biologically relevant growth factors, hormones, and inhibitors, enabling classification of modulators based on their independent, synergistic, or antagonistic effects on fusion dynamics. Consistent results obtained in an independent model further support its potential applications to additional fusion systems. By providing a robust, reproducible, and adaptable framework for time-resolved fusion analysis, this methodology bridges the gap between qualitative observation and quantitative kinetic assessment. Thus, the approach could be readily extended to other cell fusion systems following system-specific parameter optimization, offering a versatile platform for both mechanistic studies and discovery-driven screening applications.

## 1. Introduction

Cell–cell fusion is a fundamental biological process in a range of physiological and pathological conditions. Physiologically, cell fusion is essential for normal development and tissue homeostasis in diverse contexts, such as myoblast fusion during skeletal muscle formation, osteoclast differentiation in bone remodeling, trophoblast fusion during placental development, and syncytium formation during viral infection [1–7]. Dysregulated or aberrant cell fusion is implicated in cancer progression, tissue repair, inflammation, and infectious disease [8–10]. Despite its broad biological relevance, quantitative characterization of cell fusion remains technically challenging due to its dynamic, heterogeneous, and multistep nature.

Across most experimental systems, cell fusion is assessed using endpoint or semi-quantitative approaches [11] . These include immunostaining-based fusion indices, multinucleation counts, reporter gene assays, or measurement of fusion-associated gene expression and secreted factors [12–14]. While such approaches have been instrumental in identifying molecular regulators of fusion [15–19], they provide only static snapshots of a continuous process and are limited in their ability to capture fusion kinetics, intermediate states, and temporal heterogeneity within cell populations. Moreover, many commonly used assays rely on manual scoring or subjective interpretation, reducing reproducibility and scalability.

Live-cell imaging offers the possibility of directly observing fusion events in real time [20–25]. However, existing live-imaging approaches are often constrained by limited field-of-view, low throughput, or reliance on fluorescent labeling strategies that can perturb cellular behaviour [26– 29]. In addition, the lack of standardized, automated analysis pipelines has hindered the extraction of robust quantitative metrics from time-resolved imaging data [30]. A central methodological challenge is distinguishing bona fide fusion events from other processes such as cell clustering, proliferation, or migration, which can produce superficially similar morphological outcomes in label-free imaging modalities.

High-content screening (HCS) platforms combine automated microscopy with advanced image analysis and offer a powerful solution to these challenges [25,31–34]. HCS enables multiparametric, time-resolved quantification of cellular phenotypes at scale and has been widely adopted in studies of cell viability, differentiation, migration, and drug response profiling [24,33– 38]. However, despite its widespread use, there remains a lack of validated HCS-based methodologies specifically designed for real-time, label-free quantification of cell fusion dynamics. In particular, analytical frameworks that integrate segmentation, morphology-based measurements, and texture analysis to objectively capture fusion-associated cellular remodelling are not well established.

Trophoblast syncytialization is a robust and experimentally tractable model system to develop and validate such methodologies. In vitro, the human choriocarcinoma-derived BeWo cell line undergoes reproducible cell fusion in response to cyclic AMP (cAMP) signalling, most commonly induced by forskolin [39,40]. This system has been extensively used to study the molecular regulation of fusion and offers well-characterized pharmacological modulators, making it ideally suited for benchmarking quantitative fusion assays [16–19].

Here, we present a high-content, live-cell imaging pipeline for real-time assessment and quantification of cell fusion, using BeWo cell syncytialization as a model application. The method integrates automated image acquisition with a reproducible, stepwise analysis workflow combining supervised texture-based segmentation, morphology-based metrics, and intensity-independent texture analysis. We define and validate quantitative parameters that sensitively capture fusion kinetics and discriminate true fusion events from non-fusion–related cellular rearrangements. We further demonstrate the scalability of this approach by applying it to high-throughput screening of biologically relevant factors, highlighting its potential utility across diverse cell fusion contexts.

## 2. Methods

### 2.1 Cell Culture and Induction of Cell Fusion

BeWo human choriocarcinoma cells (ATCC) and HeLa cells (ATCC) were cultured in DMEM/F12 (1:1) medium (HiMedia) supplemented with 15% fetal bovine serum and 1% antibiotic– antimycotic solution (Gibco). Cells were maintained at 37 °C in a humidified incubator with 5% CO_2_ and passaged at 75–80% confluence and media were replaced every alternate day. The cells were regularly tested for mycoplasma and if positive, elimination was using commercial kit (HiMedia).

For fusion experiments, cells were seeded in 96-well glass-bottom plates (PhenoPlate, PerkinElmer) at approximately 60% confluence. BeWo cell fusion was induced by treatment with forskolin (HiMedia) (50 µM) for up to 48 h as per published protocols [40,41]. HeLa cell fusion was performed according to published protocol [42]. Briefly the cells at 60-70% confluency were treated with 50% PEG6000 (Qualigens) prepared in DMEM/F12 for 1 minute. The cells were then washed and subjected to imaging. In both the cases, control wells received vehicle alone and were maintained under identical conditions.

### 2.2 Real time PCR

Total RNA was extracted using RNA-xpress reagent (HiMedia) following the manufacturer’s protocol. RNA concentration and purity were assessed spectrophotometrically and treated with DNase I solution (1mg/ml) (HiMedia) to remove genomic DNA contamination. RNA was reverse transcribed using a high-capacity complementary DNA (cDNA) reverse transcriptase kit (Applied Biosystems) as detailed earlier [43,44]. The Bio-Rad CFX-96 thermal cycler was used to perform PCR. Each PCR reaction was run in duplicate, and gene expression was normalized to 18S rRNA, and fold change was determined [45] . The list of primers, their sequences, and optimized annealing temperatures is given in Supplementary Table 1.

### 2.3 Immunostaining

BeWo cells (5 × 10^4^ per well) were seeded in 96-well glass bottom plates and treated with forskolin (50 µM) or vehicle and incubated for 48 h. HeLa cells were seeded on glass coverslips and fusion was initiated using PEG and incubated for 48 h. F-actin staining was performed as described earlier [46]. Briefly, the cells were washed with phosphate-buffered saline (PBS) and fixed with 4% paraformaldehyde (PFA) and permeabilized in 0.5% Triton X. F-actin was visualized using fluorescently labelled CF® Dye Phalloidin Conjugates (1:200, Biotium, 00045: 20 min) then nuclei were counterstained with DAPI and mounted in hardset mounting medium (Biotium) and imaged using Operetta CLS Imaging system (Reviety).

### 2.4 Live-Cell Imaging and High-Content Image Acquisition

Label-free, real-time imaging was performed using an Operetta CLS high-content screening system (Revity) equipped with environmental control (37 °C, 5% CO_2_). Cells were imaged in digital phase contrast (DPC) and bright-field modes at hourly intervals over a 48 h period. For each well, 45 non-overlapping fields were acquired to ensure representative sampling of cellular dynamics.

### 2.5 Image Analysis Workflow and Feature Extraction

Image analysis was performed using Harmony 5.1 software (PerkinElmer) following a standardized, stepwise workflow (Fig. 1).

i. **Texture-Based Segmentation:** Initial segmentation was carried out using the PhenoLOGIC supervised machine-learning module to differentiate BeWo cell clusters from background regions based on texture features in bright-field images. Representative regions were manually annotated across approximately 20 images spanning multiple time points and experimental conditions to minimize training bias. Background regions were classified as Class A and cell clusters as Class B. The trained classifier was applied uniformly across all datasets.
ii. **Detection and Quantification of Cell Clusters:** Cell clusters were defined as contiguous regions classified as “cell” by the supervised PhenoLOGIC segmentation, where contiguity was based on pixel connectivity within segmented regions. These regions were subsequently processed using the “Find Cells” module with optimized splitting parameters to enable separation of adjacent or partially overlapping clusters while minimizing over-segmentation. Objects smaller than 1000 µm2 were excluded to remove debris and non-cellular artefacts. The total number of detected clusters per image was recorded and used for downstream analysis.
iii. **Morphological Feature Analysis:** Morphological properties of detected clusters were quantified using the “Calculate Morphology Properties” module. Parameters included total cluster area, width, length, roundness, and width-to-length ratio. Among these, the total cluster area was selected as the primary metric for assessing fusion-associated expansion.
iv. **Texture Analysis and Granularity Measurement:** Texture analysis was performed on DPC images using the “Calculate Texture Properties” module with SER (Spots, Edges, and Ridges) features. The SER Bright feature was selected as a quantitative measure of cytoplasmic granularity due to its sensitivity to fusion-associated cytoplasmic remodeling. SER features are intensity-independent owing to kernel normalization, enabling robust comparison across time points and conditions.
v. **Quantitative Metrics for Fusion Dynamics:** To capture fusion dynamics while discriminating fusion from proliferation-driven clustering, a composite metric was defined as the ratio of total cluster area to the number of detected clusters per image frame. The following formula was used.

**Figure 1.**
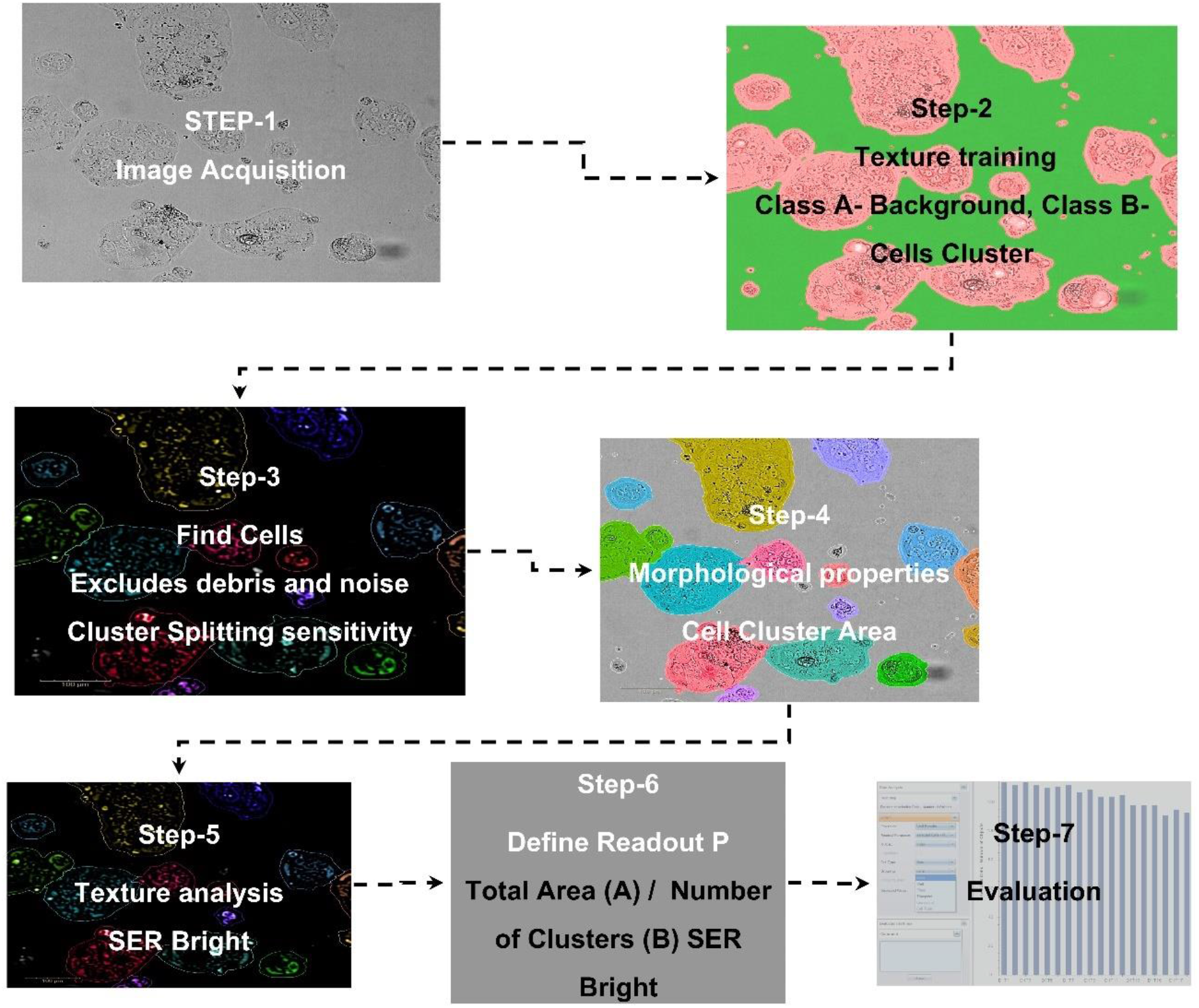
Workflow for label-free, high-content quantification of cell fusion dynamics. Stepwise image acquisition and analysis pipeline used for quantifying cell fusion dynamics. (1) Livecell image acquisition of cells using a high-content screening system over a 4811 period. (2) Supervised texture-based segmentation using the PhenoLOGIC module in Harmony 5.1 to classify background and cell cluster regions (training parameters: texture scale = 2 px, region scale = 6 px, training radius = 25 px). (3) Automated detection and splitting of cell clusters using the “Find Cells” module (minimum object area = 1000 μm^2^; splitting sensitivity = 0.75; threshold = 0.51). (4) Extraction of morphological features. (5) Texture analysis of digital phase contrast (DPC) images using SER Feature2D (scale = 4 px). (6) Calculation of the area-to-cluster ratio (total cluster area divided by number of detected clusters) and granularity as a quantitative metric of fusion dynamics. (7) Statistical analysis and data visualization.

Area-to-cluster ratio= 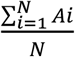 where Ai represents the area of the i-th cluster and N is the total number of detected clusters. This metric was calculated at each time point and averaged across fields and wells. Granularity values derived from SER Bright analysis were tracked over time in parallel. Together, these metrics provided complementary readouts of fusion-associated morphological expansion and cytoplasmic reorganization.

### 2.6 Validation Using Pharmacological Modulators

To assess the specificity of the analytical parameters, fusion was induced using either forskolin (50 µM) or a cell-permeable cAMP analog (1.5 µM). To inhibit fusion, we used Wortmannin, a PI3K pathway inhibitor known to block syncytialization [47,48]. BeWo cells were seeded at a density of 5 × 10^4^ cells per well in a 96-well plate, and the next day, Wortmannin (Sigma) was applied alone (100 nM) or in combination with forskolin, and imaging was done as above. Control cells received the vehicle alone. Morphological and texture-based metrics were quantified across all conditions using the same analysis workflow.

### 2.7 Effect of Various Factors on BeWo Cell Syncytialization

BeWo cells were seeded at a density of 5 × 10^4^ cells per well in a 96-well plate. The next day, cells were subjected to four different experimental conditions: (i) untreated control, (ii) 50 µM forskolin treatment, (iii) treatment with factors alone, and (iv) 50 µM forskolin along with the factor. The experimental schematic, the concentration of factors used are provided in Supplementary Figure S1 and Table S2, respectively. Working concentrations were selected based on previously published studies, as indicated by references [15,48–52]. The cells were incubated for 48 h under standard culture conditions. Live-cell imaging was performed using High-Content Screening (HCS) system, and images were acquired using the Operetta CLS Imaging system for quantitative analysis as described above.

For each condition, the slope of the area-to-cluster ratio was calculated using linear regression of the area-to-cluster ratio over the defined time window (18–42 h) corresponding to the active fusion phase to account for non-linear dynamics. Normalized values were visualized using heat maps, and hierarchical clustering was applied to classify modulators based on their effects on fusion dynamics.

### 2.8 Statistical Analysis

All quantitative data are presented as mean ± SEM. Unless otherwise stated, analyses were performed using data pooled from three independent experiments, with 10 images per condition per experiment. Statistical significance between groups was assessed using unpaired two-tailed t-tests. p-value <0.05 was considered statistically significant.

## 3 Results

### 3.1 Validation of Cell Fusion Induction in the BeWo Model System

To establish a reference system for validating quantitative fusion metrics, we first confirmed forskolin-induced cell fusion in BeWo cells using conventional molecular and morphological readouts. Following 48 h of treatment with forskolin (50 µM), cells exhibited loss of discrete cell boundaries and formation of multinucleated structures, consistent with syncytial fusion (Fig. 2A-F). Quantitative analysis revealed a marked reduction in cortical F-actin organization in forskolin-treated cells relative to controls, accompanied by a significant decrease in overall F-actin fluorescence intensity (Fig. 2G).

**Figure 2.**
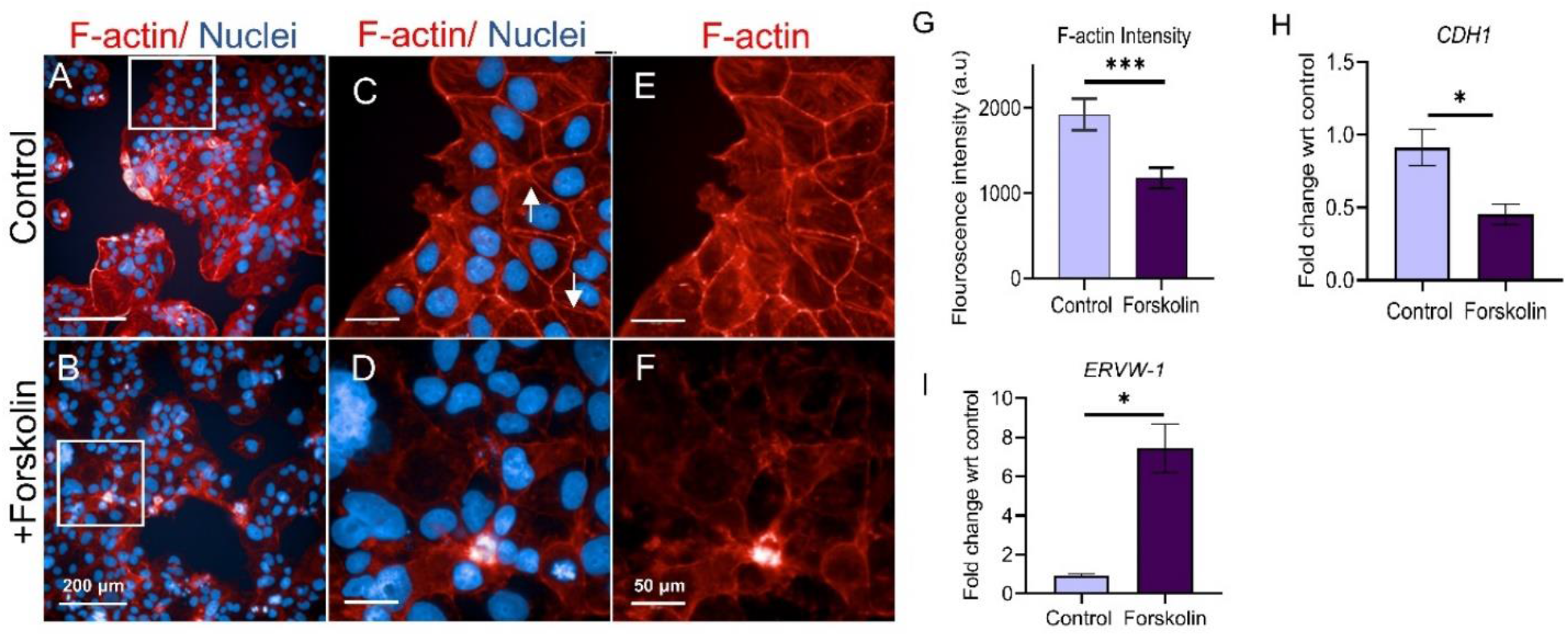
Validation of BeWo cell syncytialization using conventional molecular and imaging markers. (A-F) Representative images of BeWo cells stained fbr F-actin (Red) and nuclei (Blue). Control cells (A, C, E) exhibit discrete cell boundaries and cortical actin organization, whereas fbrskolin-treated cells (B, D, F) display loss of distinct borders and formation of multinucleated structures. Scale bars: A, B = 200 μm; C-F = 50 μm. Boxed area in A and B is magnified. (G) Quantification of F-actin fluorescence intensity in control and fbrskolin-treated cells. Data represent mean ± SEM (n = IO images per condition). Statistical significance was assessed using an unpaired two-tailed t-test (***p < 0.001).(H, 1) Relative mRNA expression of *CDH1* (H) and *ERVW-1* (Syncytin-1) (1) following forskolin treatment. Gene expression was normalized to 18S rRNA and expressed relative to untreated controls. Data represent mean ± SEM from three independent experiments. Statistical significance was assessed using an unpaired two-tailed t-test (*p < 0.05).

At the transcriptional level, expression of the adherens junction marker *CDH1* was reduced approximately twofold following forskolin treatment (Fig. 2H), while expression of the fusogenic gene *ERVW-1* (Syncytin-1) increased approximately eightfold (Fig. 2I) compared with untreated controls. Together, these observations confirmed robust induction of cell fusion and established this system as a suitable benchmark for evaluating live-cell, label-free fusion metrics.

### 3.2 Live-Cell Imaging Reveals Limitations of Area-Based Measurements Alone

To assess whether label-free live-cell imaging could resolve fusion-associated morphological changes over time, BeWo cells were imaged hourly for 48 h under control and forskolin-treated conditions. Bright-field imaging revealed progressive coalescence of cellular clusters in both groups (Fig. 3A-B) (Supplementary Video S1-S2). Although cluster merging occurred more rapidly in forskolin-treated cells, similar clustering behaviour was also observed in control cells over longer time scales, likely reflecting proliferation-driven aggregation.

**Figure 3.**
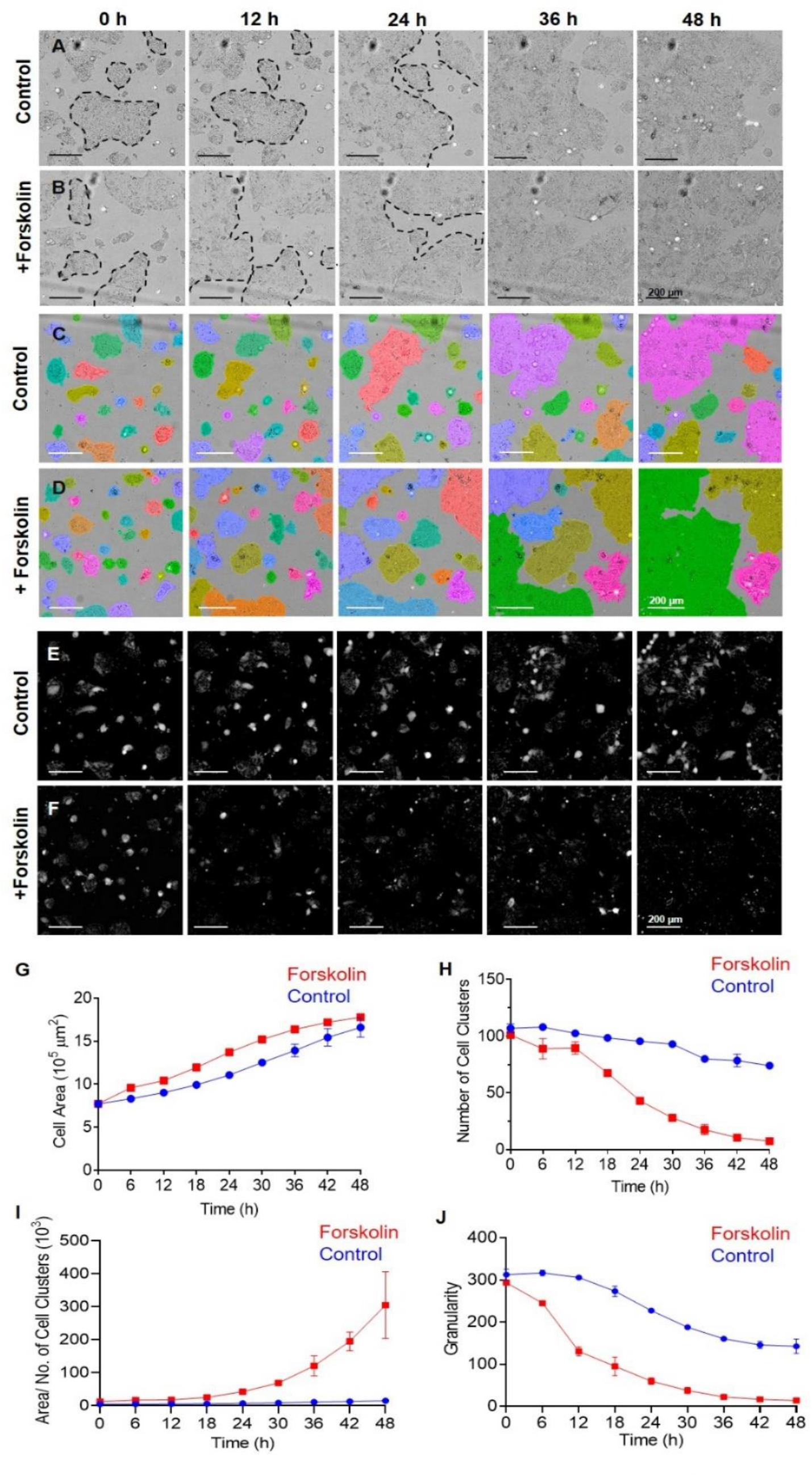
Live-cell imaging-based quantification of fusion-associated morphological and texture dynamics. Representative bright-field time-lapse images of BeWo cells cultured under control (A) or fbrskolin-treated (B) conditions over 48h. (C, D) Automated segmentation and cluster identification using Hannony 5.1 software, with system-assigned pseudocolors indicating distinct detected clusters at the indicated time points. (E, F) Representative digital phase contrast (DPC) images illustrating intracellular texture patterns in control (E) and fbrskolin-treated (F) cells. (G-J) Quantitative time-course analysis of fusion-associated parameters measured at 6 h intervals over 48h: (G) total cluster area (×10^5^ μm^2^). (H) number of detected cell clusters, (I) area-to-cluster ratio (×10^3^), and (J) cytoplasmic granularity quantified using the SER Bright feature. Control cells are shown in blue and fbrskolin-treated cells in red. Data represent mean ± SEM pooled from three independent experiments (n = 10 images per condition per experiment). Imaging was performed using an Operetta CLS high-content screening system.

Quantitative analysis of total cluster area demonstrated a gradual increase over time in both control and forskolin-treated conditions, with no statistically significant difference between groups (Fig. 3G). These observations indicate that area-based measurements alone are insufficient to discriminate true fusion events from non-fusion–related cellular growth or clustering, underscoring the need for composite or orthogonal quantitative parameters.

### 3.3 Area-to-Cluster Ratio Discriminates Fusion from Proliferation-Driven Clustering

To resolve fusion-associated morphological expansion from simple clustering, we quantified the number of discrete cell clusters at each time point using automated segmentation. In control cultures, the number of clusters decreased gradually over time (Fig. 3C). In contrast, forskolin-treated cells exhibited a more pronounced and accelerated reduction in cluster number (Fig. 3D), particularly after 18 h of treatment (Fig. 3H)

We next integrated cluster area and cluster number into a composite metric defined as the ratio of total cluster area to the number of detected clusters. This area-to-cluster ratio revealed a clear temporal divergence between conditions. While control cultures maintained relatively stable ratio values throughout the 48 h imaging period, forskolin-treated cells exhibited a marked increase beginning at approximately 24 h, consistent with the formation of large, multinucleated syncytial structures (Fig. 3I). These findings demonstrate that the area-to-cluster ratio provides a sensitive quantitative readout that distinguishes fusion-associated morphological remodeling from proliferation-driven clustering in live-cell imaging data.

### 3.4 Texture-Based Granularity Analysis Captures Fusion-Associated Cytoplasmic Remodeling

Cell fusion is accompanied not only by morphological expansion but also by intracellular cytoplasmic reorganization. To capture these changes in a label-free manner, we applied texture analysis to digital phase contrast images (Fig. 3E-F, Supplementary Video S3-S4) using the SER Bright feature, which quantifies cytoplasmic granularity independent of signal intensity.

Qualitative inspection of phase contrast images revealed a progressive reduction in intracellular texture in forskolin-treated cells relative to controls (Fig. 3J). Quantitative granularity analysis confirmed this observation: while granularity decreased modestly over time in control cultures, forskolin-treated cells exhibited an earlier onset (within 6h) and a more pronounced reduction in granularity that persisted throughout the imaging period.

Importantly, granularity measurements provided an orthogonal parameter to the area-to-cluster ratio, capturing intracellular remodeling events that are not reflected by gross morphological measurements alone. Together, these complementary metrics enabled robust, time-resolved discrimination of fusion-associated cellular states.

### 3.5 Specificity of Quantitative Metrics for Canonical Fusion Pathways

To assess the specificity of the analytical parameters for authentic fusion events, we evaluated their response to pharmacological modulation of known fusion pathways. In addition to forskolin, cell fusion was induced using a cell-permeable cyclic AMP analog. Both treatments produced comparable increases in the area-to-cluster ratio, reductions in cluster number and cytoplasmic granularity relative to untreated controls (Fig. 4A, B, and C), indicating that the metrics are responsive to cAMP/PKA-dependent fusion irrespective of the upstream stimulus.

**Figure 4.**
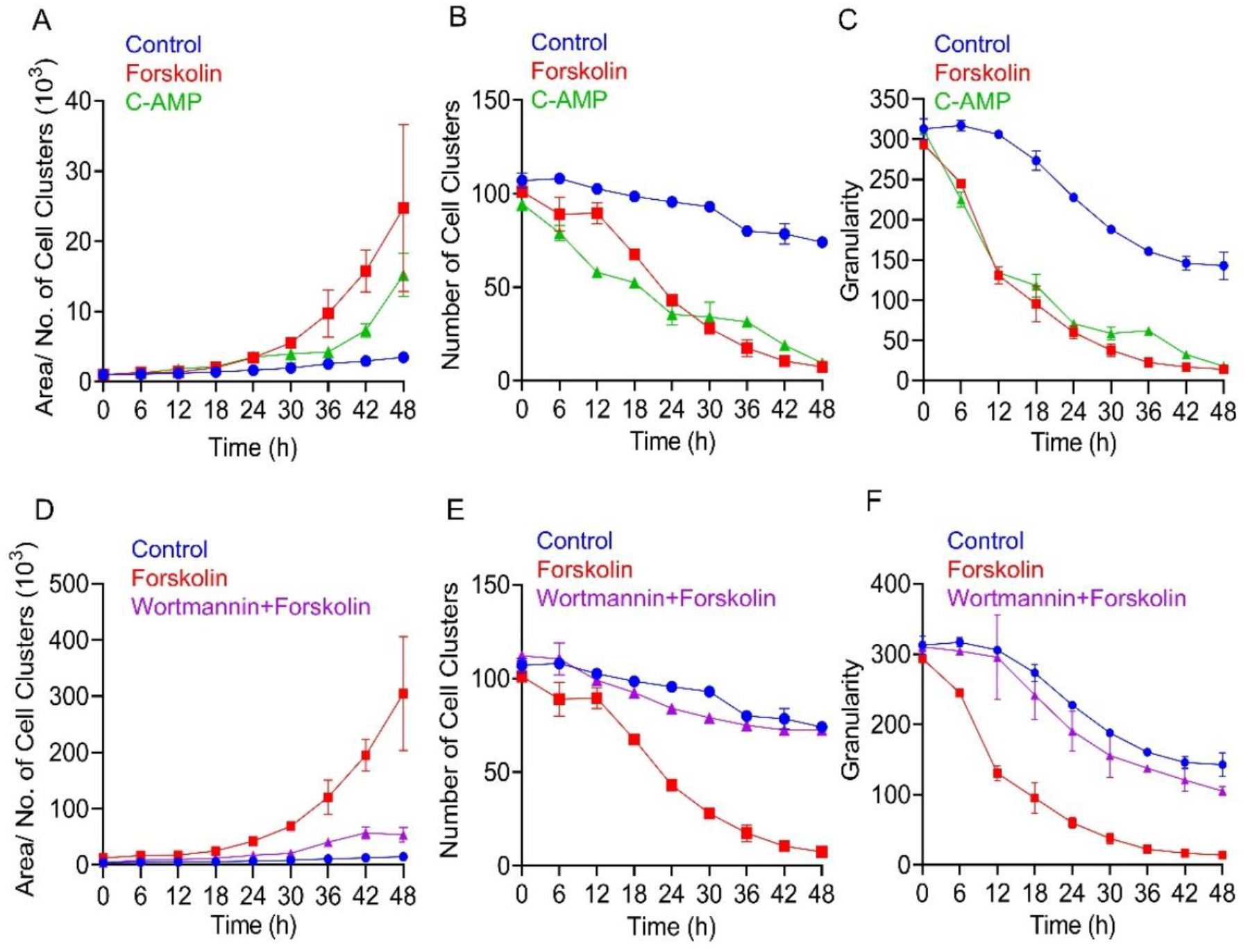
Specificity of quantitative fusion metrics using pharmacological modulators. BeWo cells were treated with forskolin (50 μM), a cell-permeable 8-Br-cAMP (cAMP, 1.5 μM), Wortmannin, or their respective combinations, and analyzed using the described imaging and analysis pipeline. (A-C) Quantification of area-to-cluster ratio (A), number of detected clusters (B), and cytoplasmic granularity (C) following treatment with forskolin or cAMP analog relative to untreated controls. (D-F) Corresponding parameters measured following treatment with forskolin alone, Wortmannin alone, or their combination. Data represent mean ± SEM from live-cell imaging experiments (n = 10 images per condition across two independent experiments). Imaging was performed using the Operetta CLS system and analysis was done using Hannony 5.1 software.

We next examined whether these parameters could detect inhibition of fusion. Co-treatment with the PI3K pathway inhibitor Wortmannin attenuated forskolin-induced changes in cluster number and granularity, yielding profiles that closely overlapped with control conditions (Fig. 4E-F). Although a modest increase in area-to-cluster ratio was observed at later time points (Fig. 4D), the absence of corresponding changes in cluster number and granularity indicated suppression of characteristic fusion-associated remodeling. These results demonstrate that the combined metric framework can distinguish partial or non-productive morphological changes from bona fide fusion events.

### 3.6 High-Throughput Screening Demonstrates Scalability and Analytical Resolution

To evaluate the scalability and utility of the pipeline for high-throughput applications, we screened fourteen biologically relevant growth factors, hormones, and inhibitors under four experimental conditions: untreated control, factor alone, forskolin alone, and factor combined with forskolin. Analysis of fusion dynamics revealed three distinct phases: an initial lag phase, a subsequent active fusion phase, and a later stationary phase (Fig. 5). Consistent with this, log transformation of the area-to-cluster ratio did not yield uniform linearization across conditions (Supplementary Fig. 2), indicating that fusion dynamics do not follow a single exponential model. Accordingly, fusion rates were quantified using the slope of the area-to-cluster ratio calculated over a restricted time window (18-42 h) corresponding to the active fusion phase.

**Figure 5.**
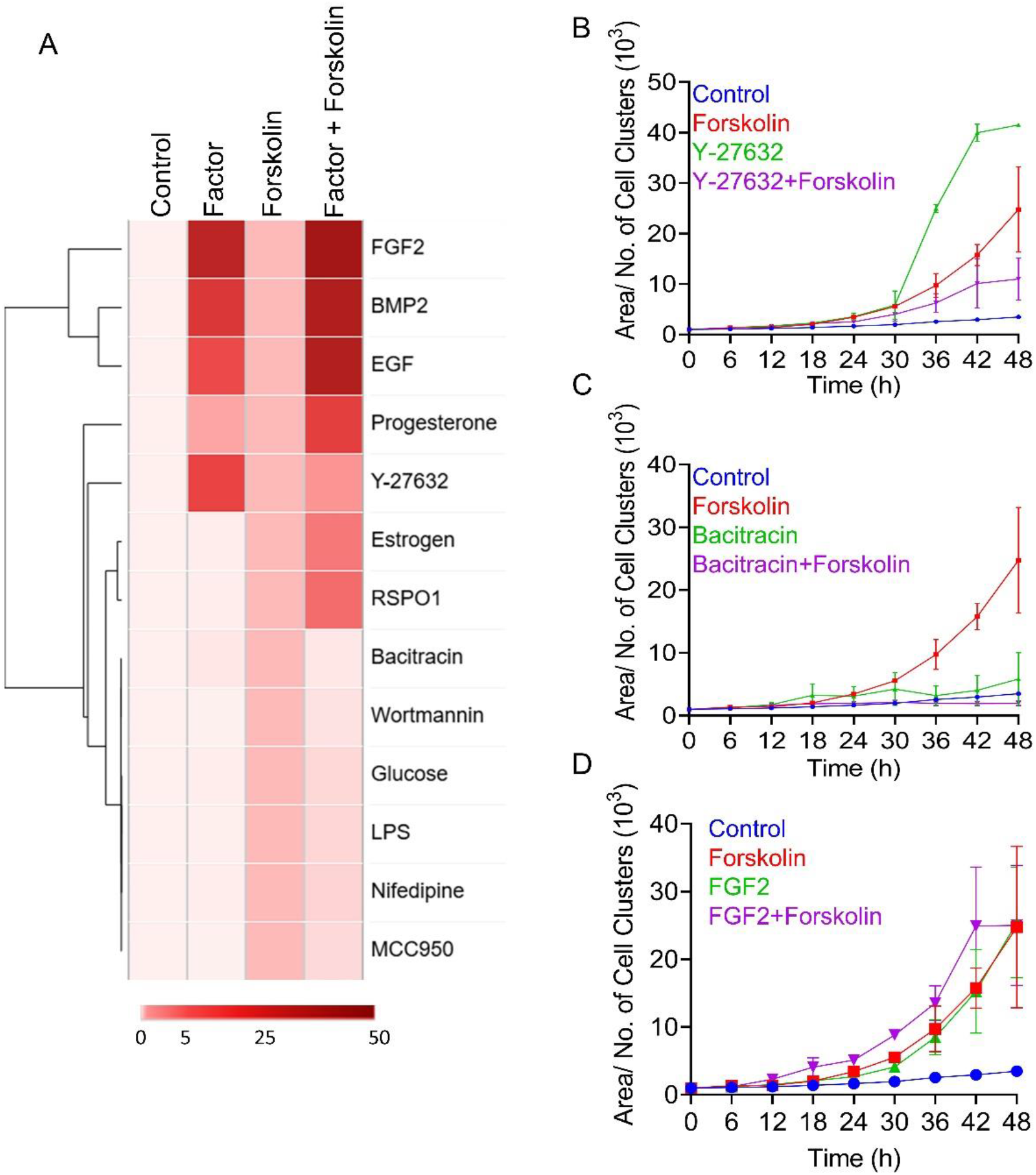
High-content screening of modulators affecting cell fusion dynamics. (A) Heatmap representation of normalized slopes (slope of fbrskolin treated cells taken as 1) of the area-to-cluster ratio over restricted time window (18-42 h) for each tested factor under four experimental conditions: untreated control, factor alone, fbrskolin alone, and factor combined with fbrskolin. Hierarchical clustering was done using Eucleadine distance. (B-D) Representative timecourse plots of area-to-cluster ratio fbr selected modulators: Y-27632 (B), Bacitracin (C), and FGF2 (D). Data represent mean ± SEM of n = 10 images per condition; two independent experiments. All imaging and analysis were performed using the Operetta CLS system and Harmony 5.1 software.

Hierarchical clustering of normalized slopes revealed distinct functional classes of modulators. Several factors promoted fusion independently of forskolin, while others exhibited synergistic or additive effects only in combination with forskolin (Fig. 5A). A subset of compounds selectively antagonized forskolin-induced fusion without exerting independent effects.

Representative examples illustrated the analytical sensitivity of the method. The ROCK inhibitor Y-27632 exhibited a biphasic response (Fig. 5B), enhancing fusion when applied alone but attenuating forskolin-induced fusion when combined. Bacitracin showed minimal activity alone but robustly suppressed forskolin-mediated fusion (Fig. 5C). In contrast, FGF2 promoted fusion both independently and synergistically with forskolin (Fig. 5D). These context-dependent effects highlight the ability of the pipeline to resolve complex, condition-specific modulation of fusion dynamics.

### 3.7 HeLa cell fusion assessment

To assess the broader applicability of the pipeline, we applied the analysis to PEG-induced homotypic fusion in HeLa cells. The pipeline successfully captured fusion-associated morphological changes, with increased area-to-cluster ratios consistent with trends observed in BeWo cells (Supplementary Fig. 3).

### 3.8 Summary of Method Performance

Collectively, these results demonstrate that the integration of morphology-based and texture-based metrics enables sensitive, specific, and scalable quantification of cell fusion dynamics using label-free live-cell imaging. The pipeline reliably distinguishes fusion from non-fusion–related clustering, captures temporal progression of fusion-associated remodelling, and supports high-throughput screening applications.

## 4 Discussion

In this study, we developed and validated a label-free, high-content live-cell imaging pipeline for quantitative analysis of cell fusion dynamics. Using trophoblast syncytialization as a model system, we demonstrate that commonly used single-parameter descriptors, such as total cell area, are insufficient to distinguish bona fide fusion events from proliferation-driven clustering in live-cell imaging datasets. By integrating cluster-based morphological expansion with texture-based granularity metrics, the proposed framework captures both structural and intracellular hallmarks of fusion in a time-resolved manner. The method reliably discriminates fusion-associated remodeling from non-fusion–related cellular rearrangements, responds appropriately to canonical pharmacological inducers and inhibitors of fusion, and scales to high-throughput screening formats capable of resolving context-dependent modulation of fusion dynamics. These findings provide a foundation for situating the approach within the broader landscape of existing fusion assays and imaging-based methodologies.

Cell-cell fusion has been studied for decades across multiple biological systems, yet its quantitative analysis has largely remained dependent on static, endpoint-based assays. Classical approaches for assessment of cell fusion include measurement of fusion indices derived from immunostaining, multinucleation counts, reporter-based assays, and expression of fusion-associated genes or hormones [10,11,53–55]. These approaches have been instrumental in identifying molecular regulators of fusion in systems ranging from myoblast differentiation to trophoblast syncytialization and virus-induced syncytium formation[2,5,10,15,19,54]. However, these methods fundamentally treat fusion as a binary outcome measured at a fixed time point, rather than as a dynamic, progressive process that unfolds over hours to days.

Several studies have incorporated live-cell imaging to visualize fusion events in real time, providing important qualitative insights into membrane remodeling and cytoskeletal reorganization during fusion [20–23,56]. Nonetheless, most live-imaging implementations rely on limited fields of view, fluorescent labeling strategies, or manual annotation, restricting throughput and reproducibility. Moreover, the absence of standardized quantitative metrics has made it difficult to distinguish bona fide fusion events from confounding processes such as cell clustering, proliferation, or migration, particularly in label-free imaging modalities. These limitations have constrained the broader adoption of live-cell fusion assays for systematic or discovery-driven applications.

In this context, the methodological contribution of the present study lies not in introducing a new biological model, but in providing a quantitative, time-resolved analytical framework that addresses these longstanding limitations. By integrating automated segmentation with morphology-based and texture-based measurements, we move beyond single-parameter readouts toward a multiparametric description of fusion-associated remodelling. Our results demonstrate that commonly used descriptors such as total cell area are insufficient to discriminate fusion from non-fusion–related cellular behaviours. In contrast, the combined use of cluster-based area expansion and cytoplasmic granularity captures both the structural and intracellular hallmarks of fusion, enabling robust discrimination of fusion dynamics in live-cell imaging datasets.

An important aspect of this framework is its reliance on image-derived features rather than molecular markers. Marker-based assays, while powerful, are inherently system-specific and often incompatible with long-term live imaging or high-throughput formats [26,27,29,57] . By contrast, morphological and texture-based features reflect downstream consequences of fusion that are shared across many fusion contexts, regardless of the upstream signalling pathways involved. Conversely, herein the inhibition experiments using Wortmannin highlight the value of multiparametric analysis in distinguishing partial or non-productive morphological changes from authentic fusion-associated remodeling.

High-content screening platforms have been widely used to study proliferation, cytotoxicity, differentiation, and migration[34,36,38], but their application to cell fusion has remained relatively underdeveloped. One reason is the lack of validated quantitative metrics tailored to fusion-specific phenotypes. By demonstrating that fusion dynamics can be captured using slope-based normalization of composite morphological parameters, our study extends the utility of high-content imaging into the domain of fusion biology. The ability to classify modulators based on their independent, synergistic, or antagonistic effects further underscores the analytical resolution of the approach and its suitability for discovery-oriented screens.

While trophoblast syncytialization was used here as a model system, this choice was motivated by experimental robustness and availability of well-characterized pharmacological modulators rather than biological specificity. The analytical principles underlying the pipeline, including cluster coalescence, area redistribution, and cytoplasmic texture remodeling, that are shared by many fusion processes, including myoblast fusion, osteoclast formation, and virus-induced syncytium formation. Indeed we could demonstrate the ability of the pipeline to detect fusion-associated changes in an independent HeLa-based system suggests that the approach is not restricted to trophoblast syncytialization and can be adapted to other fusion contexts with minimal modification. At the same time, we must emphasize that fusion systems differ substantially in cell size, geometry, kinetics, and fusion topology [24,25,33]. As such, adaptation of the pipeline to other contexts will require system-specific optimization of segmentation parameters and temporal resolution, rather than direct transfer without calibration.

Several considerations are important for interpreting and applying this approach. The pipeline quantifies fusion-associated remodeling at the population level and captures downstream morphological and cytoplasmic changes, rather than directly resolving individual membrane fusion or pore formation events. These metrics are validated against established pharmacological modulators of fusion and provide robust, time-resolved proxies of fusion dynamics. As with all image-based analyses, segmentation performance depends on image quality and may require optimization in densely packed or highly motile cultures. While the label-free design minimizes perturbation and enables longitudinal analysis[57], integration with fluorescent reporters may further enhance sensitivity in systems where fusion-associated changes are subtle or spatially restricted[24,25,57]. Collectively, these considerations reflect general features of image-based fusion assays and highlight opportunities for adaptation and extension across diverse experimental contexts.

In conclusion, this study positions high-content live-cell imaging as a practical and quantitative tool for studying cell fusion dynamics. By bridging the gap between qualitative visualization and quantitative kinetic analysis, the proposed pipeline complements existing molecular endpoint-based assays and expands the methodological repertoire available to the field. Its scalability, adaptability, and label-free design make it particularly well suited for systematic interrogation of fusion-modulating factors across diverse biological contexts.

## Supporting information

Supplementary Video_ 1_ Brightfield_Control (1)

Supplementary Video_ 2_ Brightfield_Forskolin

Supplementary Video_ 3_ DPC_Control (1)

Supplementary Video_ 4_ DPC_Forskolin

Supplementary Figure S1.JPG

Supplementary Figure S2.JPG

Supplementary Figure S3.JPG

## ACKNOWLEDGEMENTS & FUNDING

We are deeply saddened by the passing of our colleague and first author, Sandip Shinde, who contributed significantly to the conception and execution of this work. We dedicate this study to his memory.

We thank the ICMR for funding DM and the CSIR-SRF for funding AB. We are grateful to Dr. Bhakti Pathak (ICMR-NIRWoH) for kindly providing the BeWo cells and Dr Vikrant Bhor (ICMR-NIRWoH) for sharing HeLa cells. ChatGPT-5.2 assisted with language editing of the manuscript under the authors’ supervision. The manuscript bears the NIRRCH ID OA/1/12-2025

