## Supplementary figures and images for "Real-time, label-free assessment of cell fusion dynamics by high-content imaging"

### Supplementary Figure S1.JPG

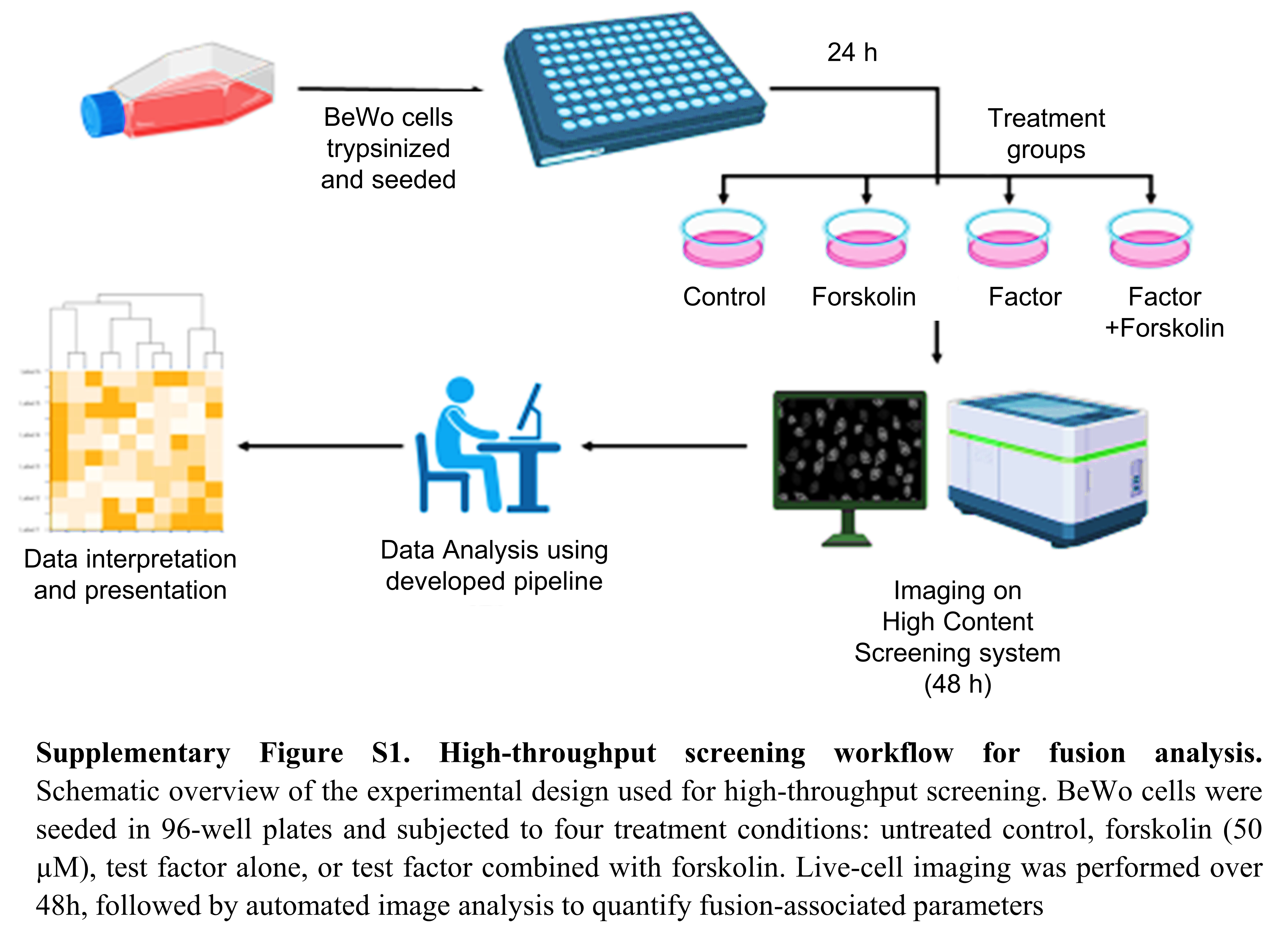

### Supplementary Figure S2.JPG

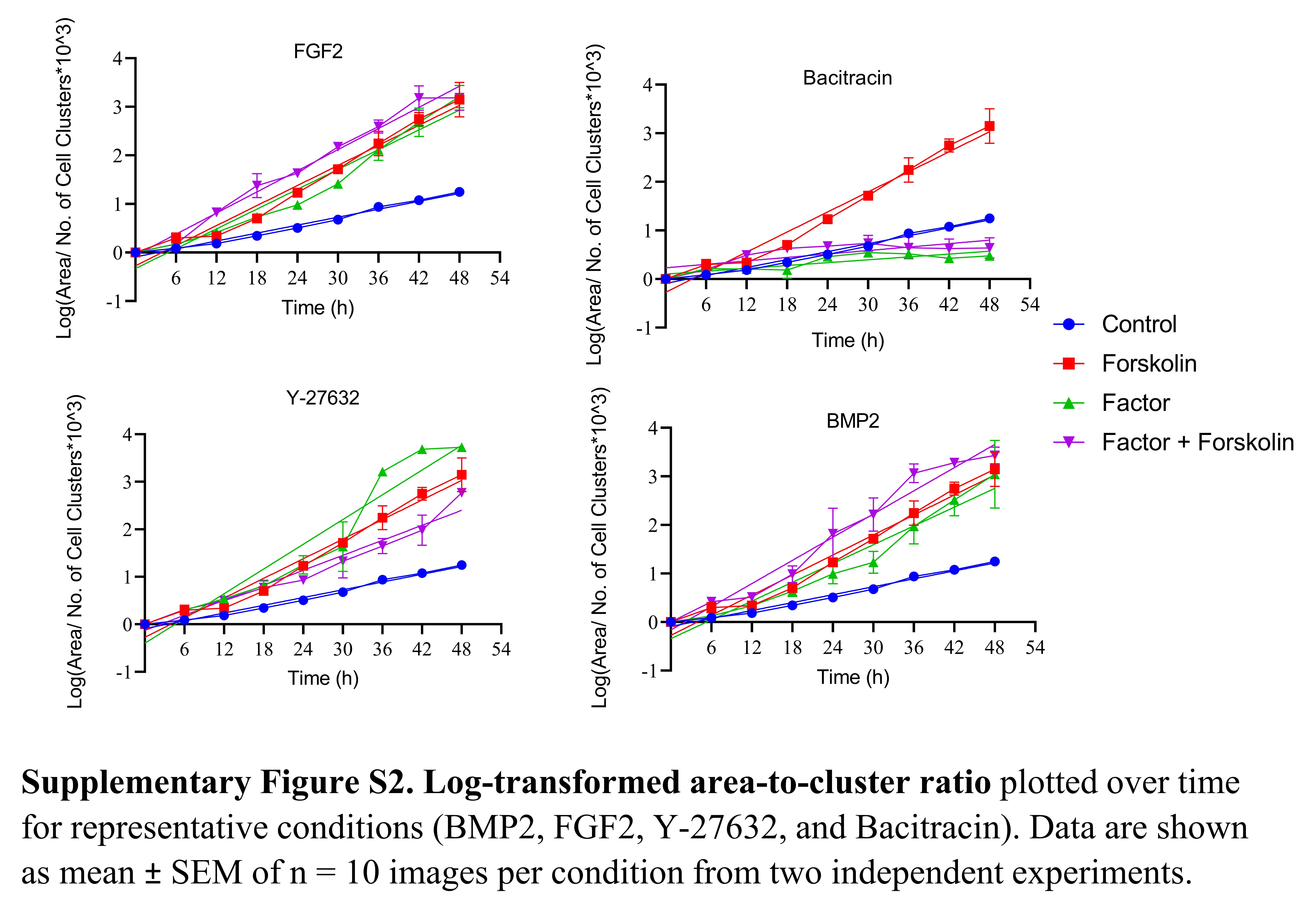

### Supplementary Figure S3.JPG

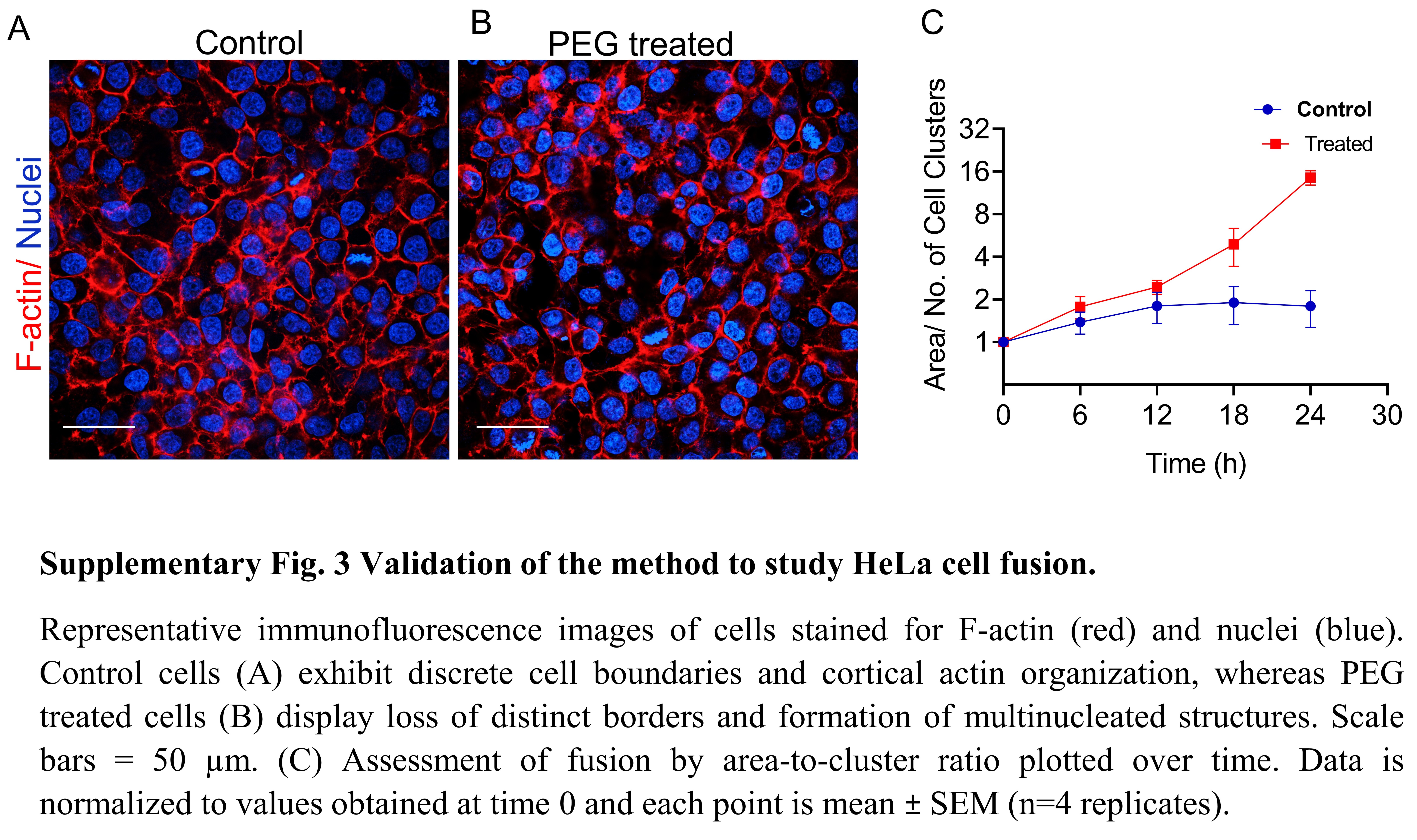
