## Supplementary Files Legends for "Real-time, label-free assessment of cell fusion dynamics by high-content imaging"

**Supplementary Data Legends


Table S2. List of Factors and Working Concentrations Used in High-Throughput Screening.**

The table lists the 15 factors tested in the high-throughput screen for their effects on BeWo cell syncytialization. Working concentrations were selected based on previously published studies as indicated by references

**Supplementary Video S1–S4**. **Live-cell imaging of trophoblast fusion.**

S1–S2: Representative bright-field time-lapse of BeWo cells under (S1) vehicle control and (S2) forskolin-treated conditions.

S3–S4: Representative digital phase contrast (DPC) time-lapse of BeWo cells (S3) vehicle control and (S4) forskolin-treated conditions.

Cells were imaged every 1h for 48h using the Operetta CLS High-Content Analysis system equipped with 37 °C, 5% CO2 environmentally controlled chamber (Revvity). The total elapsed time from the first image is indicated at the top right.
